# Comprehensive assessment of multiple biases in small RNA sequencing reveals significant differences in the performance of widely used methods

**DOI:** 10.1101/445437

**Authors:** Carrie Wright, Anandita Rajpurohit, Emily E. Burke, Courtney Williams, Leonardo Collado-Torres, Martha Kimos, Nicholas J. Brandon, Alan J. Cross, Andrew E. Jaffe, Daniel R. Weinberger, Joo Heon Shin

## Abstract

High-throughput sequencing offers advantages over other quantification methods for microRNA (miRNA), yet numerous biases make reliable quantification challenging. Previous evaluations of the biases associated with small RNA sequencing have focused on adapter ligation bias with limited evaluation of reverse transcription or amplification biases. Furthermore, evaluations of the accuracy of quantifications of isomiRs (miRNA isoforms) or the influence of starting amount on performance have been very limited and no study has yet evaluated differences in the quantification of isomiRs of altered length. In addition, no studies have yet compared the consistency of results derived from multiple moderate starting inputs. We therefore evaluated quantifications of miRNA and isomiRs using four library preparation kits, with various starting amounts, as well as quantifications following removal of duplicate reads using unique molecular identifiers (UMIs) to mitigate reverse transcription and amplification biases. All methods resulted in false isomiR detection; however, the adapter-free method tested was especially prone to false isomiR detection. We demonstrate that using UMIs improves accuracy and we provide a guide for input amounts to improve consistency. Our data show differences and limitations of current methods, thus raising concerns about the validity of quantification of miRNA and isomiRs across studies. We advocate for the use of UMIs to improve accuracy and reliability of miRNA quantifications.

## BACKGROUND

Research of miRNA expression has been instrumental in identifying miRNAs involved in development and diseases (1), and identifying expression-signatures for use as biomarkers (2–4). Small RNA sequencing (sRNA-seq) allows for detection of novel miRNAs and altered canonical miRNA sequences, termed isomiRs (5–7). These miRNA isoforms are produced by shifts in Drosha and Dicer cleavage sites of the pri- and pre- miRNA sequence, as well as trimming by exoribonucleases, additions of bases by nucleotidyl transferases, and RNA editing by adenosine deaminase acting on RNA (ADAR) enzymes (6). isomiRs show differences in stability, localization, and functionality. Some isoforms can even regulate alternative repertoires of mRNAs as compared to the canonical sequence (6, 8–12). Recent research indicates that they are of clinical importance for many diseases and conditions, including cancer (13), diabetes (14), and Huntington’s disease (15). Despite the enhanced capability to detect such sequences, there has been little assessment of the quantification of isomiRs using sRNA-seq; and miRNA quantifications using this method are often inconsistent across studies (16). This is likely in part due to differences between methods and/or variation in the detection by individual methods (17) (from library preparation to preprocessing to normalization, etc.). Furthermore, several aspects of sRNA-seq can lead to the preferential quantification of some miRNAs and reduced or completely lacking quantification of others, thus introducing biases that lead to misrepresentations of true miRNA expression levels (18). Evaluations and comparisons of the accuracy (how close measurements are to the truth) and consistency (how close measurements are across replicates) associated with current methods are critical for proper cross-study interpretation and for guiding methodological improvement.

Evidence suggests that biases and inconsistencies in sRNA-seq based quantifications and group comparisons are largely based on study design and library preparation methods (17). The details of these issues have been reviewed elsewhere (16, 19–25). Some of these issues are avoidable with proper study design. However, bias and inconsistency related to adapter ligation, cDNA synthesis, and amplification may principally be dependent on library preparation and preprocessing methods, which are less readily controlled.

A considerable number of studies have evaluated adapter ligation bias in quantifications from several commercially available kits (24, 26–28); however, to our knowledge, only one study has directly compared the performance of randomized adapter methods and adapter ligation-free methods (24). Furthermore, limited studies have investigated the influence of reverse transcription or amplification bias in sRNA-seq (29–32) and no study to date has evaluated the use of unique molecular identifiers (UMIs) in order to identify and remove duplicate reads to mitigate such biases in sRNA-seq biological samples. While amplification bias is associated with variations in sequence length and GC content (33), and while the use of UMIs has proven to be useful in mitigating amplification bias for traditional RNA sequencing (34, 35) and has become increasingly popular, the exact implementation of this method is still unresolved in the RNA sequencing field at large (36). It remained unclear if UMIs would also be useful for small RNA sequencing as this has not been explored to date. The consensus in the field about PCR amplification bias in small RNA has been divided. One view holds that amplification bias appears to be minimal in sRNA-seq. They argue this because small RNA sequences are very similar in size, studies show that bias in sRNA-seq largely appears to be due to adapter ligation bias (18, 37, 38), and studies show no distortion of quantification results with excessive numbers of PCR cycles compared to more reasonable numbers of cycles (31, 32). However, others suggest that amplification bias in sRNA-seq could also introduce bias as it does in traditional RNA sequencing (10), especially given the large range of GC content among miRNAs. While a couple of studies have used UMIs in sRNA-seq (29, 30), only one has evaluated the reproducibility of sRNA-seq quantifications obtained from utilizing UMIs to those without (29), in which the authors concluded that biological technical replicates had less variation when UMIs were used to remove duplicate reads compared to when either all or no duplicates were removed. However, no statistical tests were performed in this assessment, and no evaluation of the influence of the UMI deduplication on the accuracy of the quantifications was performed. In the sRNA-seq literature there has also been little assessment of the influence of starting amount on the consistency of quantifications. While RNA editing detection has been evaluated (26), other aspects of isomiR quantification have not yet been performed.

To complete the gap left by previous studies, we comprehensively evaluated bias among miRNA and isomiR quantifications from four commercially available library preparation methods, as well as those obtained following the removal of duplicate reads using UMIs. We also evaluated the consistency of the results using a variety of starting amounts. We assessed the similarity of the quantifications from each method, the diversity of the detection of different types of small RNAs by each method, as well as the accuracy and the consistency of the results obtained from each method within and across batch. Such evaluations are critical for optimizing sRNA-seq methods to obtain both reliably consistent and accurate results across batches and studies, and to therefore allow for more accurate and reproducible miRNA and isomiR quantifications in disease states and conditions. Based on these results, we offer suggestions for future study designs.

## RESULTS

### Study Design

In this study we evaluated the influence of several potential sources of bias and inconsistency on miRNA quantifications (**Fig. 1.a**) by comparing the performance of four commercially available kits including two that are designed to mitigate adapter ligation bias in different ways (**Fig. 1.b**) and two preprocessing methods including one to mitigate reverse transcription and adapter ligation bias and a control for comparison (**Fig. 1.c**), as well as various starting amounts (100ng to 2,000ng) for each method to determine the reliability of results achieved with smaller inputs (**Fig. 1.d**). The following library preparation kits were compared: 1) the Clontech SMARTer smRNA-Seq Kit for Illumina, now owned by Takara Bio (Clontech), which incorporates adapter and index sequences during reverse transcription and amplification and is therefore ligase-free to avoid adapter ligation bias; 2) the Bioo Scientific NEXTflex Illumina Small RNA Sequencing Kit v3 (NEXTflex), now owned by Perkin Elmer and called NEXTFLEX, which utilizes adapter sequences with random nucleotide sequences adjacent to the miRNA binding location giving each miRNA a variety of adapter sequences to bind to avoid adapter ligation bias; 3) the Illumina TruSeq Small RNA Library Prep Kit (Illumina); and 4) the New England BioLabs Next Multiplex small RNA kit (NEB). Based on our literature search Illumina and NEB sRNA-seq kits appear to be the first and second most widely used kits to date, respectively. The NEB and the NEXTflex kits include polyethylene glycol in an effort to reduce adapter ligation bias by improving overall ligation efficiency.

**Fig. 1.**
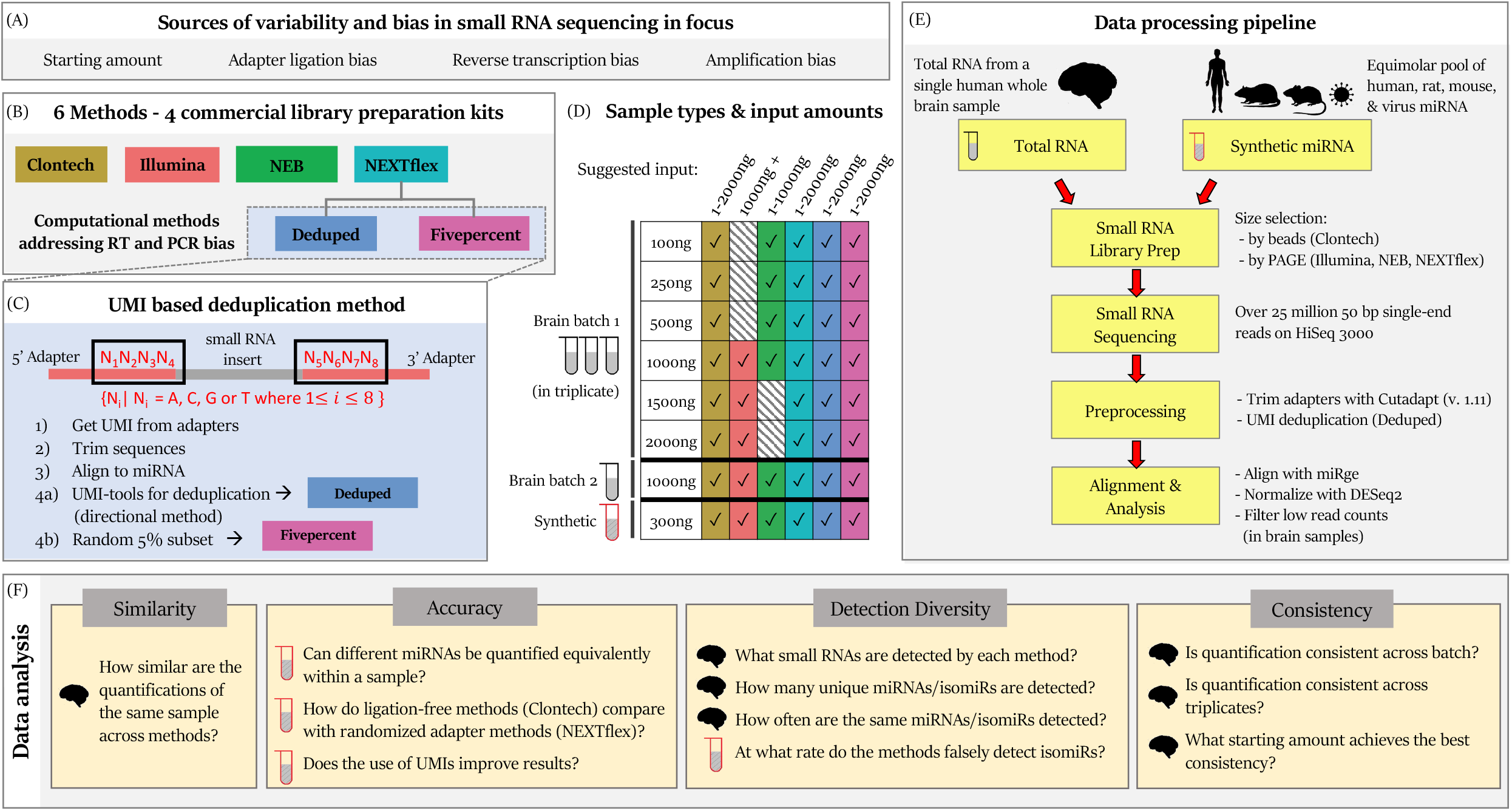
Study Design. **(a)** We evaluated the influence of starting amount on the consistency of results, as well as the accuracy of results obtained when using a variety of methods, including those intended to reduce bias from adapter ligation, reverse transcription (RT), and PCR amplification. **(b)** We compared four commercially available kits and two preprocessing methods to address RT and PCR bias. **(c)** In the Deduped method we collapsed duplicate reads based on a unique molecular identifier (UMI) of degenerate bases in the adapter sequences (bases within the black boxes). We also compared the collapsed data with a 5% subset of the NEXTflex data to determine if performance differences were due to the UMI-based collapsing of reads or simply due to having fewer reads. **(d)** We evaluate two data types: miRNA quantifications from homogenate whole brain total RNA and miRNA quantifications from a pool of 962 equimolar synthetic RNAs with sequences corresponding to human, rat, mouse, and virus miRNA. We had two batches of human brain data. The first included triplicates of different starting amounts based on the kit manufacturers’ suggested ranges. The second included a single sample of the same human brain with 1000ng of input. We used 300ng of the synthetic miRNAs for each tested method. **(e)** Our processing pipelines for the two types of RNA studied. **(f)** We evaluated the 6 small-RNA sequencing methods using 4 major assessments. The brain icon indicates utilization of brain samples to assess a question, while the red tube indicates utilization of synthetic miRNA samples.

We also evaluated the influence of reverse transcription and amplification bias by utilizing the random sequences within the adapters of the NEXTflex kit (that are added prior to the cDNA synthesis and PCR amplification steps), as UMIs. These UMIs allow for the removal of duplicate reads introduced during amplification and possible mitigation for sequences that may have been preferentially reverse transcribed (**Fig. 1.c**). We will hereinafter refer to these data as “Deduped”. To determine if differences identified between the Deduped data and the NEXTflex data were simply due to a reduction in the number of reads (as the UMI-based deduplication process reduces the data down to 5% of the original), we also included a random 5 % subset of the NEXTflex reads, hereinafter referred to as “Fivepercent,” for comparison.

We evaluated two types of samples (**Fig. 1.d-e**) and processed the data following the methods outlined in **Fig. 1.e**. See the methods for more details of our experimental approach. We then evaluated several questions shown in **Fig. 1.f** about the similarity of the quantifications obtained from the 6 tested methods (**Fig. 1.b**), the accuracy of those quantifications using synthetic miRNAs in equimolar concentration, the ability of each method to detect a variety of miRNAs and isomiRs, and the consistency of the quantifications by each method of technical replicates within the same batch and across two batches.

### Similarity – Overall quantifications are similar, yet results for individual miRNAs are quite divergent across methods

We first performed a general evaluation of the similarity of the resulting miRNA quantifications from each method (**Fig. 1.b**) and of major contributors to overall variability using the data derived from the same human brain sample across technical replicates (**Fig. 1.f.Similarity**). Hierarchical cluster analysis indicated that the samples generally clustered by method and starting amount (**Fig. 2.a**). Differential analysis of the miRNA expression estimates revealed that the methods (**Fig. 1.b**) produce overall relatively similar results, however some individual miRNAs showed very different quantifications with intensity ratios ranging as extreme as −9 to 6 (**Fig. 2.b**).

**Fig. 2.**
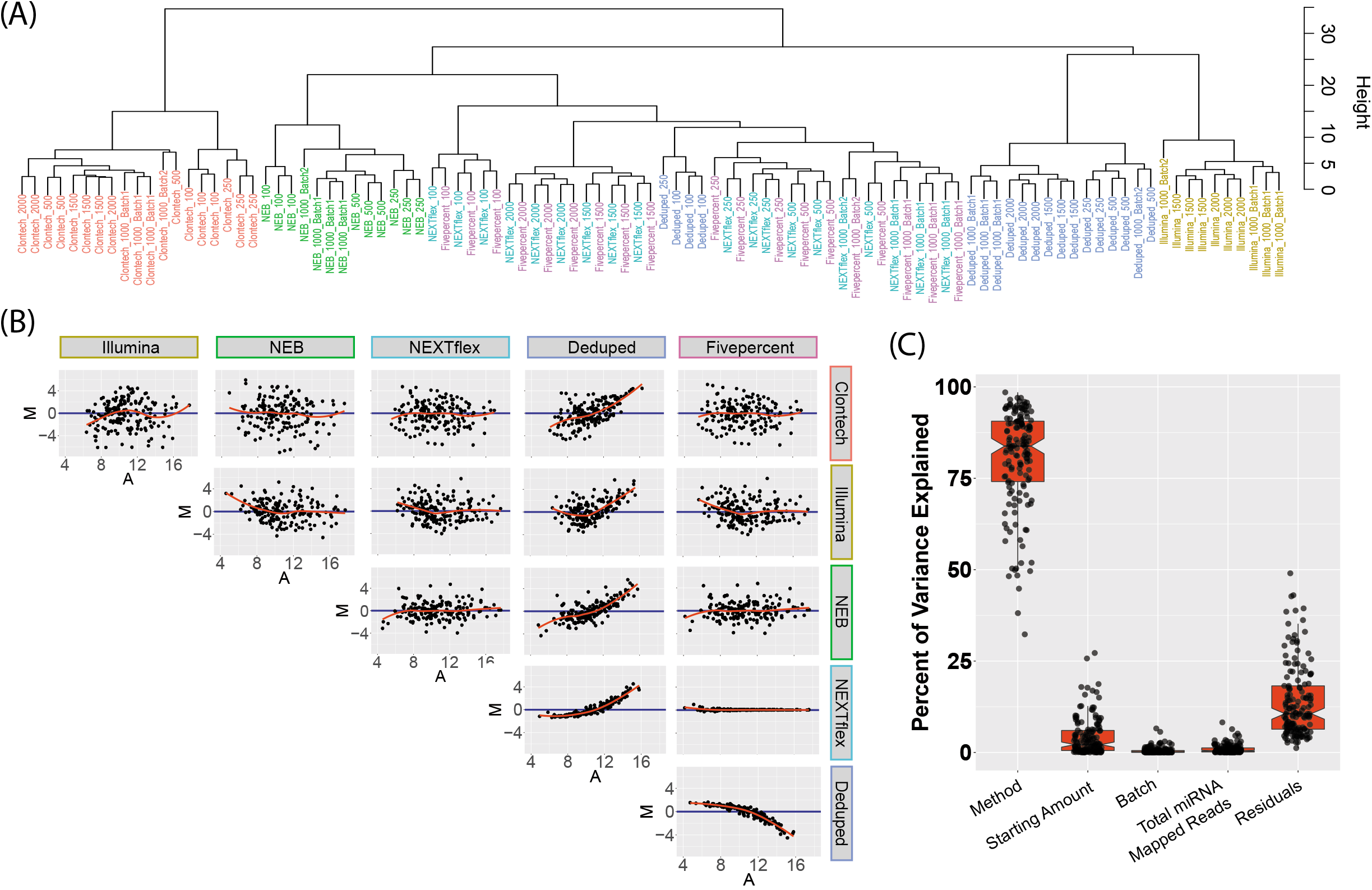
Similarity Assessment. **(a)** Dendrogram depicting cluster analysis shows that samples largely cluster by method and starting amount. **(b)** Individual points represent the miRNAs quantified by all of the methods; the y-axis of each plot shows the log ratio of the normalized quantification estimates between the two methods, while the x-axis shows the average expression. These plots are referred to as MA plots. **(c)** The percent of variance explained by method, starting amount, batch, the number of reads mapped to miRNA, and the variance unaccounted for by these factors. Each point represents the variance explained by each factor for an individual miRNA sequence that was quantified by all of the tested methods.

Evaluating the top 20 abundant miRNAs from each method (**Supplementary Table 1**), only 6 miRNAs (30%) overlapped across all methods (however the top 20 for Fivepercent were identical to the top 20 from the raw full NEXTflex data). Thus, emphasizing only the most abundant miRNAs for further study may be problematic. The overlap between the most abundant miRNAs detected by Clontech and the other methods was lower (45% to 55%) than the overlap between Illumina, NEB, and NEXTflex (60-65%). The Deduped method resulted in an 85% overlap with the raw NEXTflex data.

Sum of squares analysis revealed that method choice was the largest contributor to miRNA count variability (on average 82% variance explained for individual miRNAs (**Fig. 2.c**) when evaluating the data from all methods (excluding the Fivepercent control). This further exemplifies the lack of consistency in quantifications that may occur when different methods are utilized.

### Accuracy – Reduction of numerous biases improves accuracy

To assess the accuracy of each method (**Fig. 1.f.Accuracy**), we investigated how consistently each kit detected 962 equimolar synthetic miRNA sequences. We calculated the difference of each miRNA count from the mean count for all miRNAs for each method, which we called “accuracy error”. The six methods showed significant differences in accuracy (F = 40.00, p < 2.2e-16). The Deduped data had significantly less accuracy error compared to all other methods (up to ≈ 8% less error), followed by comparable accuracy for Clontech, NEXTflex, and Fivepercent methods (which did not significantly differ from one another in post hoc analysis), and worse accuracy for the NEB and Illumina methods. This suggests that the Illumina and NEB methods detect different sequences with less validity than the other methods. This was expected, given the known adapter ligation bias associated with these methods. Our results suggest that the methods utilized by the Clontech and NEXTflex kits both diminish bias – likely due to a reduction in adapter ligation bias. Using UMI sequences for deduping resulted in additional error reduction (the raw NEXTflex data had 2.81% more error) (**Fig. 3.a and Supplementary Table 2**), which may be due to a reduction in reverse transcription and/or amplification bias. This is consistent with our analysis of the overall variance of the counts for these synthetic sequences (**Fig. 3.b**). The concordance of the rank of the sequences with higher accuracy error across the methods was poor (data not shown), suggesting that different sequences were prone to bias for each of the methods.

**Fig. 3.**
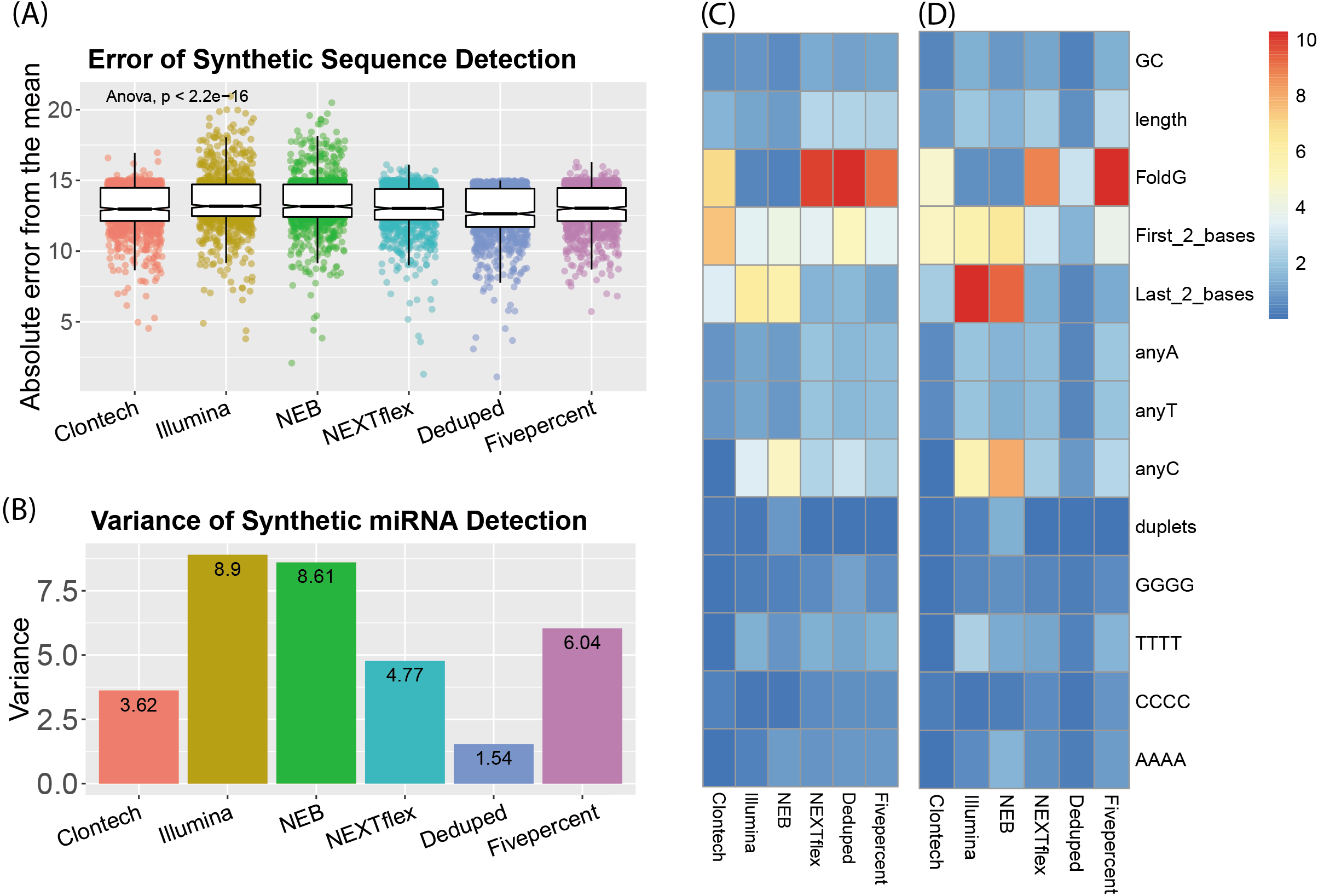
Accuracy Assessment. **(a)** Individual points represent the absolute difference of each synthetic miRNA quantification from the mean of all quantifications of the equimolar synthetic sequences for each small RNA sequencing method. **(b)** The variance of all the quantification estimates for the synthetic sequences. **(c)** The percent variance of synthetic sequence quantifications explained by each of these sequence characteristics: GC content, length, Gibbs free energy of the predicted secondary structure (FoldG), identity of the first (5’) and last (3’) two bases, the count of individual bases, and the presence of repeat sequences, such as duplets of the same base or quadruplets of the same base. The heatmap legend shows the percentage of variance from 0 to 10 percent. **(d)** The percent variance explained by each of the sequence characteristics but weighted by the overall variance of each method, as shown in B. The heatmap legend shows the percentage of variance from 0 to 10 percent.

We thus analyzed the overall contribution of different sequence characteristics to the variance of the count estimates of the synthetic miRNAs and found that indeed different characteristics were associated with variability for the different methods (**Fig. 3.c, Fig. 3.d**). The secondary structure Gibbs free energy was highly influential for Clontech (explaining ≈ 7% of the variance), and the NEXTflex-based methods (explaining ≈ 10 % of the variance for each). The identity of the last 2 bases was influential for all methods but in particular for the NEB and Illumina methods (explaining ≈ 6 % of the variance for each), suggesting that adapter ligation of the 3’ end particularly introduces bias of miRNA quantifications, in agreement with previous work (32).

The identity of the first 2 bases (5’) was most influential for Clontech and explained ≈ 8% of the variance suggesting that the SMART template-switching of the 5’ end may introduce more bias. The number of Cs within a sequence also accounted for a relatively large percentage of the variance (2.5-5% for all methods except for Clontech). Interestingly, GC content only accounted for ≈ 1% of the variance for each method. See **Supplementary Fig. 1** for more detailed information.

### Detection of RNA classes - Libraries generated using the Clontech Kit had very low miRNA mapping rates

We next assessed the percentage of reads that mapped to miRNA or other small RNA species for each of our brain-derived samples using *bowtie* (39) (**Fig. 1.f. Detection Diversity**). We excluded the Deduped data and its control, as alignment was required to produce these data. There was a significant difference in the miRNA mapping rate of the 1000ng starting input data across the kits (F = 108.9, p-value = 5.73e-09). The NEXTflex and NEB methods had the highest rates, while the Clontech method had the lowest mapping rate, with only 1-2% of all reads mapping to miRNAs (**Fig. 4.a-b**), as previously described (24) (**Supplementary Table 3**). There was a significant difference for all the tested types of RNA across the methods except for small Cajal body-specific RNAs (scaRNA) after multiple testing correction. The Clontech reads largely aligned to ribosomal RNA (rRNA) and had significantly higher rates of small nucleolar RNAs (snoRNA) and small nuclear (snRNA) mapping than the other methods, while the NEXTflex method resulted in the largest number of P-element induced wimpy testis (PIWI)– interacting RNA (piRNA) reads (**Supplementary Table 4**). All of the kits had quite consistent mapping rates across the various starting amounts (**Fig. 4.b**). Mapping rates of the synthetic RNA were much more comparable among the methods, suggesting that the differences seen with the biological samples are largely due to differences in detection of other biological RNAs (**Supplementary Table 3**).

**Fig. 4.**
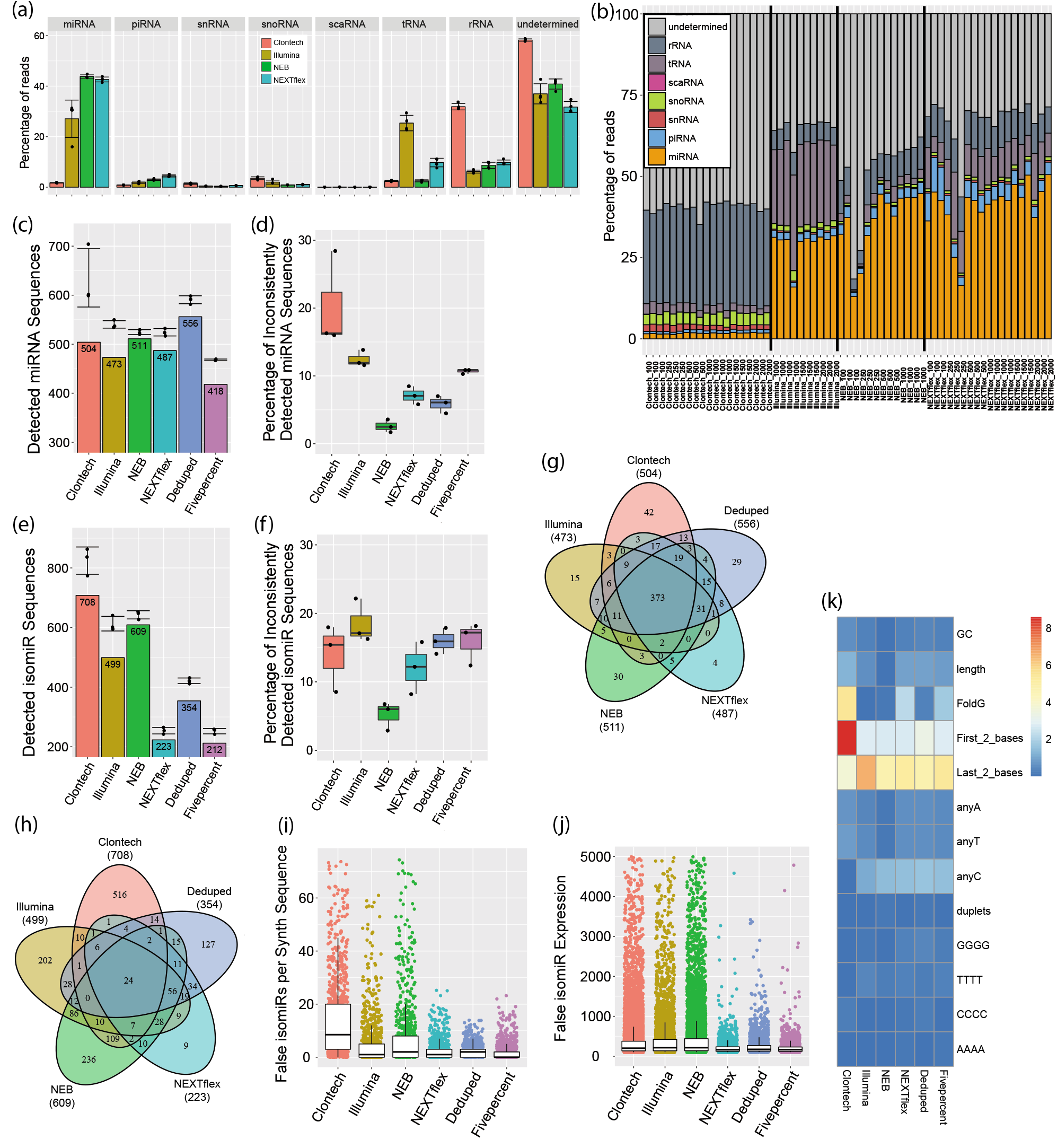
Detection Diversity Assessment. **(a)** Mapping rate of various small RNAs utilizing the 1000ng input human brain data. Undetermined indicates that the read did not map to the annotations of the evaluated small RNA classes. Error bars show standard deviation. **(b)** Mapping rate of small RNAs for all starting input amounts for each method. The Y-axis shows the percentage of reads of each category and the X-axis shows each tested brain sample. **(c)** The bars show the number of unique miRNAs with greater than 10 normalized reads common to all triplicates for the 1000ng data of the first batch. Points indicate the number of unique miRNAs for each triplicate and the standard deviation error bars shown are for these points. **(d)** Percentage of miRNAs that had quantifications above 10 in only 1 or 2 of the triplicates. **(e)** Bars show number of unique isomiRs with greater than 100 normalized reads common to all triplicates for the 1000ng data of the first batch. Points indicate the number of unique isomiRs for each triplicate and the standard devation error bars shown are for these points. **(f)** Percentage of isomiRs that had quantifications above 100 in only 1 or 2 of the triplicates. **(g)** Overlap of unique miRNAs with greater than 10 normalized reads in all triplicates for the 1000ng data of the first batch. **(h)** Overlap of unique isomiRs with greater than 100 normalized reads in all triplicates for the 1000ng data of the first batch. **(i)** Number of false isomiRs detected for each of the 962 synthetic sequences. **(j)** Number of normalized reads (expression) of the false isomiRs. **(k)** Percent variance of the number of isomiRs observed for each synthetic sequence explained by various sequence characteristics. The heatmap legend shows the percentage of variance from 0 to 9 percent.

### Detection of unique miRNAs - The Deduped data and Clontech data had better detection rates

To discern if any of the methods have an advantage in detecting a diversity of unique miRNAs, we compared the detection rate of miRNA sequences (**Fig 1.f. Detection Diversity**). Here we define a miRNA as detected if the miRNA was present with at least 10 normalized reads in the quantifications for each of the triplicates of the 1000ng batch 1 brain data. The number of detected unique miRNAs was highest in the Deduped data, and lowest in the Fivepercent and Illumina data (**Fig. 4.c**), which was consistent when including the second batch (**Supplementary Table 5**). Despite the low mapping rate of the Clontech samples, the miRNA diversity detected by this kit was relatively comparable to that of the other methods tested. Since both the Deduped and the Fivepercent data also included only 5% of the total raw NEXTflex reads, both of these methods also resulted in a much lower number of reads that could map to miRNA. The similarity of the detection rates of all the methods despite the large difference in miRNA mapping rates is due to the DESeq2 (40) normalization strategy utilized, which accounts for differences in library composition, and the high sequencing depth. An analysis of subsamples containing only 10 million, 5 million or 1 million reads of the Clontech data resulted in lower detection diversity (**Supplementary Table 6**).

Using the data from all starting amounts, there was a significant difference in the number of detected miRNAs across methods (F = 7.69, p-value = 0.00017), however pairwise comparisons were largely nonsignificant (except between Clontech and Fivepercent, p = 0.0024). There was a weak but significant positive relationship (r = 0.4, p-value = 0.027) between detection diversity and input amount (**Supplementary Fig. 2**) using all methods. Thus, as anticipated, larger inputs resulted in a more diverse pool of detected unique miRNAs; however, the pool size did not differ greatly. When evaluating each kit individually, only the Deduped method had a significant (p-value = 0.009) and strongly positive relationship between starting amount and the number of unique detected miRNAs (r = 0.92).

### Detection consistency - The Clontech method was significantly worse than the others

To determine how well each method consistently detected the same miRNAs (**Fig. 1.f. Detection Diversity**), we calculated the proportion of miRNAs detected for each sample that were not detected by the other two samples within the triplicates as a measure of detection inconsistency using the 1000ng input data (**Fig. 4.d**). There was a significant difference in the inconsistency of detection overall between the methods (F = 12.27, p-value = 0.0002), and although no individual contrasts between pairs of methods were significant in post-hoc analysis, the Clontech data resulted in the highest level of inconsistency, and NEB performed the best, with the lowest level of inconsistency.

Analysis of the full set of data including all starting amounts (**Supplementary Fig. 2**) demonstrated a significant difference in the inconsistency across methods (F = 14.83, p-value = 1.65e-10) and starting amounts (t = −3, p-value = 0.00257, Pearson *r* = −0.31). There was significantly more inconsistency for the Clontech method compared to all other methods (up to 240%) except the Fivepercent control method (**Supplementary Table 7**). This suggests that although the Clontech level of detection may have been rescued by the high depth of sequencing, the low mapping rate may still result in much poorer consistency of detection.

### Detection of isomiRs - The methods detected significantly different numbers of isomiRs – the Clontech method detected the most

We next evaluated the isomiR detection rate of each method (**Fig. 1.f. Detection Diversity**). We define an isomiR as detected if it had greater than 100 normalized reads in all triplicates for each method of the 1000ng input data. We observed the largest number of unique isomiR sequences in the Clontech data and the lowest in the NEXTflex data (**Fig. 4.e**). When evaluating detection across both batches, the Clontech data remained the most diverse (with the greatest number of isomiRs consistently detected in both batches), while the Illumina detected the lowest number of unique isomiRs (**Supplementary Table 5**). Using all the data derived from all the starting amounts, we determined that there was a significant difference in the number of isomiRs detected across the methods (F = 83.5, p-value = 8.89e-15), but not across starting amounts (**Supplementary Fig. 2**). Clontech detected the largest number (up to 250% more), followed by NEB (up to 169% more) and Illumina (up to 147% more), while the NEXTflex based methods similarly detected the least (**Supplementary Table 8**).

When evaluating the consistency of isomiR detection (**Fig. 4.f, Supplementary Fig. 2**), there was a significant difference in the triplicate consistency of detection (F = 5.9, p-value = 0.006), but again no individual contrasts between pairs of methods were significant. For the 1000ng input data, the Illumina data had the highest inconsistency, while the NEB data had the least.

### Detection overlap - Despite different miRNA mapping rates, all the methods capture overlapping miRNAs but very few overlapping isomiRs

We next characterized the overlap of unique miRNA sequences captured by each method (**Fig. 1.f. Detection Diversity**). Evaluating the miRNAs consistently detected by all 1000ng triplicates of the first batch, we determined that in general a large proportion of the miRNAs were commonly detected by all of the methods (on average 74%), and only a small fraction of miRNAs was uniquely detected by a single method (4.7% on average) (**Fig. 4.g**). In contrast, on average only 5% of the detected isomiRs by each method overlapped those detected by all the other methods (**Fig. 4.h**).

### Detection of false positive isomiRs - All methods detected false isomiRs, especially Clontech and NEB

To assess the possibility that the methods may result in false isomiR detections, we evaluated the presence of isomiRs in the synthetic miRNA data (which has no designed isomiRs) (**Fig. 1.f. Detection Diversity**). False isomiRs were detected by all of the tested methods. We compared the methods based on: 1) the number of overall unique detected isomiR sequences, 2) the number of unique isomiR sequences detected for each individual synthetic sequence, and 3) the quantifications of the individual false isomiRs. There was a significant difference in the number of isomiRs detected for each synthetic sequence by the methods (F = 176.37, p-value = <2.2e-16). The Clontech method detected more unique isomiRs overall than all the other tested methods (on average 401% more false isomiRs); 14,130 were observed for Clontech, 5,021 for Illumina, 9,074 for NEB, 2,221 for NEXTflex, 2,213 for Deduped, and 1,779 for Fivepercent. The number of unique isomiRs observed for each synthetic sequence was also significantly higher. On average the Clontech method resulted in 14 isomiRs per synthetic sequence, while the NEXTflex-based methods (the raw NEXTflex data, the Deduped, and the Fivepercent) resulted in roughly 2 isomiRs per synthetic sequence (**Fig. 4.i, Supplementary Table 9**). The counts observed for the individual isomiRs detected were significantly higher for NEB than all the other methods (with 60% higher expression than the isomiRs detected by the NEXTflex based methods) (**Fig. 4.j**). The NEXTflex methods (raw, Deduped, and Fivepercent) resulted in the fewest isomiRs detected, with the fewest isomiR counts per synthetic sequences, and with the lowest expression. There was no difference between the Deduped and the raw NEXTflex data for the expression of the isomiRs or in the number detected per synthetic sequence (**Supplementary Table 9**).

Sequence feature analysis revealed that the identity of the first two bases (5’) accounted for most of the variance in the number of isomiRs detected for each synthetic sequence for the Clontech kit (accounting for nearly 9% of the variance) (**Fig. 4.k**). Therefore, false positive isomiRs may be generated during the reverse transcription step of the library preparation for this method. This is consistent with other studies that suggest that the template-switching reverse transcription method utilized by Clontech of the 5’ end can lead to shortened miRNA transcripts in a process called strand invasion (41) and potentially longer miRNA transcripts due to concatamers of the template-switching oligo (42). In contrast, the last 2 bases (3’) accounted for the largest amount of variance of the other methods (on average 5.3%).

### Consistency across Batch - Illumina had the lowest consistency, while the other methods performed similarly

To determine the consistency of results obtained across batches for each method, we compared the mean of the triplicates in one batch to a second batch of a single sample of the same human brain (**Fig. 1.f.Consistency**). Using the normalized and log transformed reads for miRNAs that were found to be detected by each kit when filtering for greater than 10 reads across all 4 samples for each kit, we calculated the distance from the mean of the two batches for each detected miRNA individually. Overall, method choice had a weak significant association with error across batch (F = 2.39, p-value =0.036). This association was driven by the batch error of the Illumina method which was significantly higher than the other methods, with up to 74% more error than other methods (**Fig. 5.a, Supplementary Table 10**). NEB, NEXTflex, and the Deduped data were the most consistent across batch, with no significant difference in the performance of these methods (p>0.05). The top miRNAs showed some level of concordance across the methods (data not shown).

**Fig. 5.**
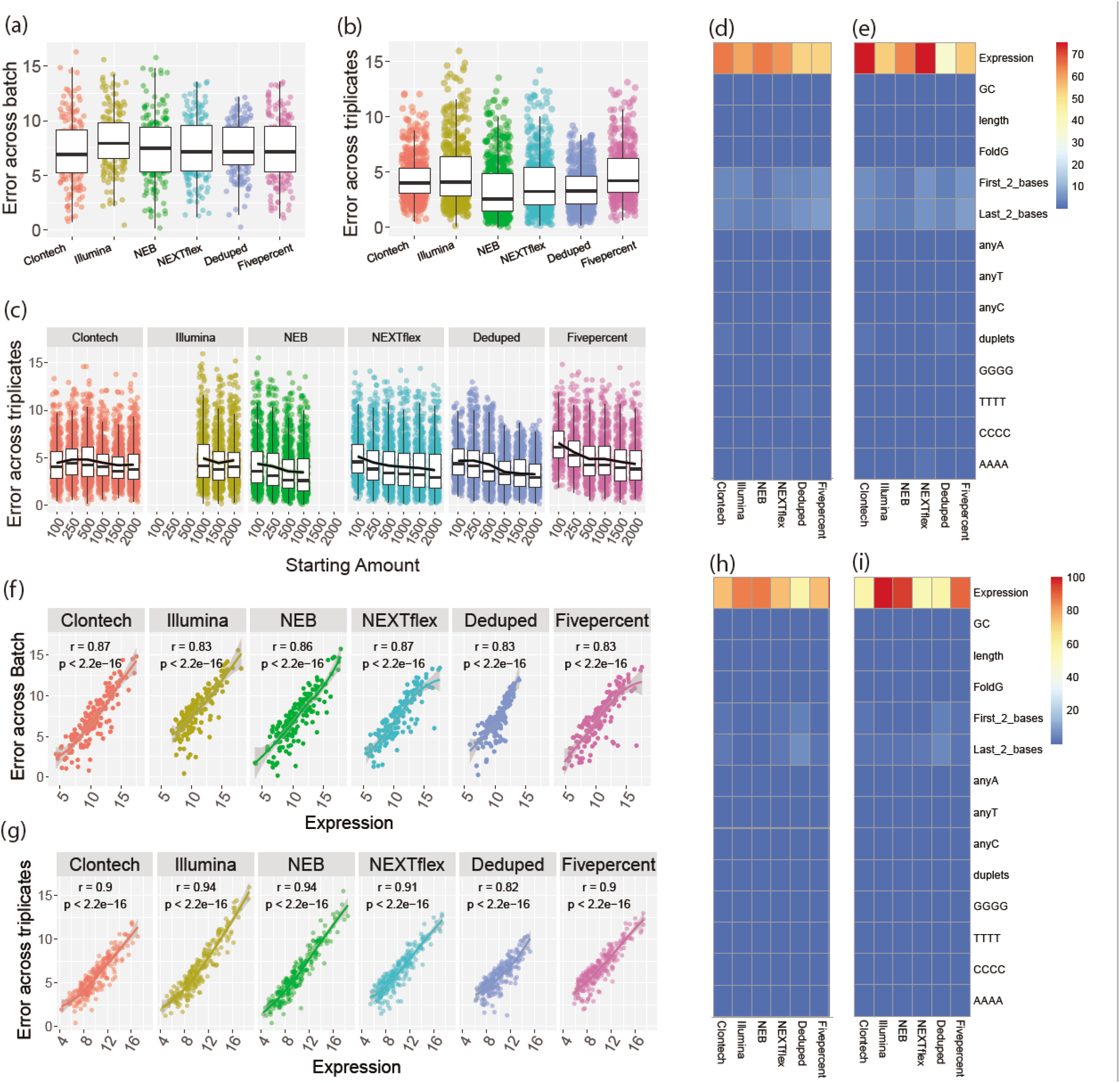
Consistency Assessment. **(a)** Absolute difference of the normalized and log_2_ transformed quantifications (norm_quantifications) of the second batch from the mean of the triplicates of the first batch for each quantified miRNA of the 1000ng input data. **(b)** Absolute difference of norm_quantifications for each quantified miRNA from a given triplicate to that of the mean of all three triplicates of the 1000ng input data. **(c)** Absolute difference of norm_quantifications for each quantified miRNA from a given triplicate to that of the mean of all three triplicates of the data for all the starting inputs. **(d)** Percent variance of batch inconsistency (a) explained by various sequence factors. The heatmap legend shows the percentage of variance from 0 to 75 percent. **(e)** Percent variance of batch inconsistency (a) explained by various sequence factors weighted by the overall batch variance of each method. The heatmap legend shows the percentage of variance from 0 to 75 percent. **(f)** Plots of the association of expression and batch error. **(g)** Percent variance explained by various sequence factors of the triplicate inconsistency plotted in **(c)**. The heatmap legend shows the percentage of variance from 0 to 100 percent. **(h)** Percent variance explained by various sequence factors of the triplicate inconsistency plotted in C and weighted by the overall variance of triplicate error for each method. **(i)** Plots of the association of expression and triplicate inconsistency using all starting input data in **(c)**. The heatmap legend shows the percentage of variance from 0 to 100 percent.

### Consistency across triplicates - Clontech and Illumina had the lowest consistency

We then evaluated the consistency of the triplicates (**Fig. 1.f.Consistency**) within the 1000ng data, by calculating the distance of each triplicate from the mean of all three triplicates. We determined that there was a significant difference across the methods (F = 36.7, p-value = <2.2e-16). Consistency was significantly higher for the NEB and Deduped methods, while Clontech, Illumina, and the random Fivepercent had the lowest consistency (with ≈ 20-40% more error, **Fig. 5.b, Supplementary Table 11**). Deduping of the NEXTflex data improved consistency. The raw data had 14% more error between triplicates.

We then calculated the triplicate consistency for each starting amount. We determined that using all starting amount data, there was still a significant difference in triplicate consistency between the methods (F = 79.7, p-value =<2.2e-16), but there was no relationship with starting amount (Pearson *r* = −0.11) (**Fig. 5.c**). All pairs of methods were significantly different, except for the contrasts between NEB and Deduped and between Clontech and Illumina (**Fig. 5.c, Supplementary Table 11**). NEB and Deduped again had the greatest consistency (up to 23% less inconsistency) and Clontech and Illumina had the least (≈17% more inconsistency).

### Factors affecting consistency - The most abundant miRNAs were the most inconsistently detected for each method

To determine if any aspect of the miRNA sequences was associated with more or less consistency across batch (**Fig. 1.f.Consistency**), we evaluated the association of various sequence factors with the batch error estimate. For each method, the expression of each individual miRNA was the largest contributor to variance of batch error (**Fig. 5.d-e**). All methods showed a significant and positive relationship between expression and inconsistency across batch (Pearson *r* >= 0.83 for all methods), **Fig. 5.f**.

Evaluating sequence characteristic associations with triplicate consistency, again, expression was the largest contributor to variance of error estimates (**Fig. 5.g-h**) and again all methods showed a significant positive association with expression and inconsistency across triplicates (Pearson >0.82 for all methods), **Fig. 5.i**.

To determine if the same miRNAs showed high error across the starting amounts or methods, we ranked the triplicate consistency error estimates and compared the ranks between the starting amounts of a given method and between methods (data not shown). The concordance of the ranks between starting amounts and methods was highest among the sequences with the highest error with roughly 40% concordance.

### Consistency and its relationship to starting amount - There was no improvement in consistency beyond 500ng of total RNA for most methods

Using data normalized and filtered for greater than 10 normalized reads for each method individually, we further evaluated the influence of starting amount on consistency across triplicates (**Fig. 1.f.Consistency**). Overall, starting amount was significantly associated (p<0.05) with triplicate consistency error for each method except for Illumina, which is likely due to the fact that fewer starting amounts were assessed for this method, **Fig. 5.c, Supplementary Table 12**. The results suggest that a larger starting amount will generally improve consistency, see **Supplementary Table 12** for specific guidance for each kit. For most methods the highest consistency with the lowest starting amount was achieved with 500ng, however, 1000ng improved the consistency of the Deduped data. The consistency was relatively similar for all the Clontech kit samples regardless of starting amount.

## DISCUSSION

We report an extensive evaluation of commonly used sRNA-seq kits for their performance in identifying and quantifying miRNAs and isomiRs, as well as the results obtained with the use of a UMI and a UMI control. Our detailed analyses identify critical factors that influence their performance. Prior performance evaluations of current sRNA-seq methods have been very limited and adapter ligation bias has largely been the focus of earlier reports (26, 31, 38, 43). Several studies have compared the NEXTflex kit with the Illumina and NEB kits (24, 26–28, 44, 45), and most suggest that the NEXTflex kit offers advantages due to reduced adapter ligation bias by including randomized adapters. We compared the NEXTflex kit with the Clontech kit which was also designed to mitigate adapter ligation bias, but by using an adapter ligation-free method. Only one prior study has compared the performance of these two kits using a previous and now discontinued version of the NEXTflex kit (24), which demonstrated that the Clontech kit resulted in less bias, however, only 6 synthetic miRNAs were utilized in their accuracy assessment. A recent study performed at the same time as ours agreed with our findings that these two kits perform similarly for accuracy (46). A similar UMI method is utilized by a recent library preparation kit by Qiagen. However, this kit was released after the data collection of our analysis. In addition, this kit, similarly to the NEB and Illumina kits, does not include methods to reduce adapter ligation bias, and the UMI is added after reverse transcription, which therefore does not allow for any reduction in bias associated with this step. The results of a recent study, which performed a similar analysis as ours, further suggest that the Qiagen kit has more bias and is less accurate than the Clontech and NEXTflex Kits (46).

To better assess the contributions of bias in the quantifications resulting from various library preparation designs, we have evaluated the quantifications from each method using a variety of metrics including: 1) Similarity – how similar are the quantifications across methods (**Fig. 1.f.Similarity**) ; 2) Accuracy – how well does each method equally quantify different equimolar miRNAs (**Fig. 1.f.Accuracy**); 3) Detection diversity – what capacity does each method have to capture a diverse range of unique small RNAs (**Fig. 1.f. Detection Diversity**); and 4) Consistency – how similar are results across batches, triplicates, and different starting inputs (**Fig. 1.f.Consistency**). Our analysis of individual sequences using the metric tests provide information about potential bias due to adapter ligation, reverse transcription, and amplification. Table 1 summarizes our results. Overall, there are clear and important differences between the methods tested and all show performance limitations in real world sRNA-seq. Based on our results, we propose a number of suggestions for future studies.

**Table 1.** Summary of Results

| Category | Assessment | Clontech | Illumina | NEB | NEXTflex | Deduped | Fivepercent |
| --- | --- | --- | --- | --- | --- | --- | --- |
|  | Adapter ligation mitigation? | Adapter Ligation free | None | PEG | PEG and Randomized adapters | PEG and Randomized adapters | PEG and Randomized adapters |
|  | RT and PCR bias mitigation? | None | None | None | None | UMI | None |
| **Performance Metrics** |  |  |  |  |  |  |  |
| **Similarity** | How similar are the quantifications of the same brain sample? | lowest correlations with other methods | Intermediate | Intermediate | Intermediate | Intermediate | Intermediate |
| **Accuracy** | Accuracy - similar detection of different sequences | Intermediate | Worst | Worst | Intermediate | Best | Intermediate |
| **Detection Diversity** | Mapping rate to miRNA | Worst | Best | Best | Best | Best* | Best* |
|  | Detection of miRNA across triplicates at 1000ng | Intermediate | Intermediate | Intermediate | Intermediate | Best | Worst |
|  | Inconsistency of miRNA detection at 1000ng | Worst | Intermediate | Best | Intermediate | Intermediate | Intermediate |
|  | Detection of isomiRs across triplicates at 1000ng | Best | Intermediate | Intermediate | Worst | Intermediate | Worst |
|  | Inconsistency of isomiR detection at 1000ng | Intermediate | Worst | Best | Intermediate | Intermediate | Intermediate |
|  | False IsomiR detection per synthetic sequence | Worst | Intermediate | Intermediate | Best | Best | Best |
|  | False IsomiR expression | Intermediate | Intermediate | Worst | Best | Best | Best |
| **Consistency** | Consistency across batch | Intermediate | Worst | Best | Best | Best | Intermediate |
|  | Consistency across triplicates | Intermediate | Worst | Best | Intermediate | Best | Worst |
| **Conclusion** |  |  |  |  |  |  |  |
| **Accuracy** | Accuracy | Intermediate | Worst | Worst | Intermediate | Best | Intermediate |

Table 1. Legend. The table depicts the results of the various assessments performed on the 6 small RNA sequencing methods. *The mapping rate of the raw NEXTflex data should reflect the quality for the data used for the Deduped and Fivepercent methods as these are derived from the raw NEXTflex data.

### Similarity

First, cross-study comparisons using different methods should be viewed with skepticism, because although the kits resulted in fairly similar results overall, quantifications of individual miRNAs, including the most abundant miRNAs, varied widely across methods (**Supplementary Table 1, Fig. 2.b**). More research is required to determine how to best utilize data derived from different sRNA-seq methods for mega- and meta-analyses. We also advise against further study of only the top expressed miRNAs from a single sRNA-seq study, particularly when a more biased method is utilized, as the top observed miRNAs may not be truly among the most abundant or influential, but instead those that are preferentially observed by the method. This issue has previously been discussed at length^16^. It is important to note that it remains unclear how relatively abundant a miRNA needs to be to exert biological importance in different contexts.

### Accuracy

We suggest the use of a degenerate base method, such as NEXTflex or a ligation-free method, to improve accuracy. These methods appeared to equally improve accuracy, likely due to a reduction of adapter ligation bias (**Fig. 3.a-b**). We suggest that future small RNA studies utilize a UMI strategy, as our NEXTflex data preprocessed to account for UMI duplicates was even more accurate, reducing the overall variance of the log_2_ transformed and normalized quantifications by 68%, or on average the difference from the mean for each miRNA by nearly 3% (**Fig. 3.a-b, Supplementary Table 2**). We speculate that our deduplication method led to such improvements due to reduced reverse transcription and/or amplification bias. Our sequence-specific analysis further indicated that secondary structure of miRNAs was one of the largest contributors to error of the Clontech and NEXTflex kits for the accuracy assessment, which appeared to be mitigated in the UMI deduped data for the NEXTflex kit (**Fig. 3.c-d**). This suggests that the secondary structure of miRNA sequences may be particularly influential for reverse transcription and/or amplification bias, in agreement with previous work that indicates that secondary structure can indeed influence reverse transcription (47). More work is required to determine the extent that amplification or reverse transcription are particularly contributing to bias, and to what extent each are mitigated by the use of UMIs. Furthermore, it is unclear if the use of deduplication would improve other methods beyond the performance level of the current NEXTflex kit. However, the UMIs are inherently already included in the NEXTflex adapters, making this one of the best current options to mitigate bias in sRNA-seq.

### Detection diversity

All of the methods tested were capable of detecting a diverse range of miRNA sequences and there was a high degree of overlap in the identity of the miRNAs detected by each method (**Fig. 4.c-e**). Therefore, any of the tested methods may be appropriate for assessments about general miRNA diversity. However, the identity of miRNAs detected by Illumina varied greatly across batch, **Supplementary Table 5**). We observed greater resolution for detection of a larger variety of miRNAs with greater sequencing depth. We did not evaluate depths above 20 million reads, so it remains unknown if even greater resolution can be achieved beyond this depth, however subsets of our 20 million depth data resulted in a reduction of diversity. We also observed that a more diverse pool was detected with larger input amounts; therefore, for the best diversity of detection, we recommend using the largest input possible.

The Clontech kit resulted in the largest percentage of reads mapping to snoRNAs and snRNAs, while the NEXTflex kit resulted in the largest percentage of piRNA mapping reads (nearly 4.2 times higher than Clontech) (**Fig. 4.a-b, Supplementary Table 4**). Therefore, if these particular small RNAs are of interest, we would suggest the use of these two kits respectively. We did not evaluate the diversity of these other classes of small RNAs; however, given the results of our miRNA analysis, we predict that deeper sequencing will result in greater resolution of diversity.

We especially suggest using randomized adapter methods, such as NEXTflex, for studies involving isomiR analysis. We suggest that all isomiR studies utilize an additional method for validation, as all methods resulted in the observation of false isomiRs. We acknowledge that some observed isomiRs in the synthetic data may be due to errors in the synthesis of the synthetic miRNA pool, however the differences between the methods suggests that some methods may detect more false isomiR sequences due to improper adapters or other features of the library preparation. In particular, the Clontech method resulted in the highest level, thus we do not suggest that others utilize this method for studies that aim to evaluate isomiR expression (**Fig. 4.h-j**). Furthermore, because this method utilizes polyadenylation of the 3’ end, it is impossible to truly distinguish isomiRs that terminate with 3’ adenine bases. We speculate that some of the false isomiRs detected in the other methods may also result from PCR stutter, in which sequences with repeats or low entropy may experience deletions or expansions during PCR amplification (36, 48). In all, the Deduped method resulted in the highest number of detected miRNAs with the lowest false isomiR detection (**Fig. 4.h-j**). Therefore, of the tested methods, we suggest that the Deduped method may be the best for detecting the most diverse and reliable set of miRNAs. Further work is necessary to determine the potential sources that result in the detection of these false isomiRs and how this may influence the quantification of isomiRs in biological samples.

### Consistency

The Deduped method was also the most consistent for individual miRNA quantifications across triplicates within the same batch (**Fig. 5.b**) Therefore, we suggest the use of this method for optimal consistency. In general, we particularly caution against the use of Illumina when multiple batches of sequencing will be involved in a study, as this method resulted in significantly more inconsistent results across batches relative to all the other tested methods (**Fig. 5.a, Supplementary Table 5**)

An earlier analysis determined that detection consistency was poorer with much smaller starting amounts (10ng) (27). Agreeing with this, our results indicate that larger starting amounts for some methods may mitigate inconsistent quantifications of miRNAs and isomiRs. Overall, we observed the most consistent results across triplicates when utilizing 500ng or greater of starting input. In most cases, 500ng was sufficient, and no improvement was achieved with higher input amounts. However, the Deduped method performed best with at least 1000ng and the Clontech method resulted in similar levels of consistency despite the use of smaller inputs (**Supplementary Table 12**). Thus, if differing starting amounts or smaller starting amounts are required, and interest in isomiRs is limited, the Clontech method may be the best choice.

Additional studies of UMI use for other library preparation methods and across biological samples are necessary to further understand the ability of UMIs to improve the consistency and reproducibility of sRNA-seq studies. Further work is also necessary to optimize the length of the UMI. In some cases, all UMIs will become saturated if a given small RNA is very highly expressed. Our calculations suggest that this UMI length is sufficient for the brain (using our current methods), in which miRNA make up a very small fraction of the total RNA and in which our data suggested that the most abundant miRNA represented only 11% of the miRNA reads. However longer UMIs may be required for tissues with greater enrichment of miRNAs or greater enrichment of other small RNAs of interest, where single RNAs may have more than 65,536 individual copies before amplification (see **Supplementary Note 1**, which refers to **Supplementary Table 13**).

## CONCLUSIONS

In conclusion, we observed significant differences in the accuracy, detection, and consistency of the various sRNA-seq methods tested suggesting that the methods differ in terms of bias contributed by adapter ligation, reverse transcription, and amplification. Our results underscore the importance of the library preparation methods and suggest that with moderately large starting amounts, the NEXTflex kit with deduplication may produce the least biased and most consistent results within and across studies. Our results suggest that bias is introduced in sRNA-seq due to reverse transcription and/or amplification and that the use of UMIs should be considered for further optimization to mitigate these biases in future sRNA-seq studies. Additional work is needed to decipher the role of these biases in sRNA-seq in order to guide more accurate sequencing methods. Further, and perhaps most noteworthy, our results indicate that all methods may result in false isomiR detection, and that the Clontech template switching method results in the detection of substantially more false isomiRs. Therefore, we advise caution with isomiR quantifications particularly when using this method. Ultimately, additional standardization of sRNA-seq data generation and analysis will improve our ability to understand the expression and regulatory role of these small but important RNAs in conditions and disease.

## METHODS

### Library preparation and sequencing

Two sample types were used to evaluate the performance of the sRNA-seq methods, (**Fig. 1.d**). To evaluate detection and consistency we used various starting amounts in triplicate of total RNA from a homogenate human brain sample, purchased from Ambion and derived from a 74-year-old Caucasian female. The cause of death of this individual was respiratory failure. To evaluate accuracy, we used 300ng of the Miltenyi Biotec miRXplore Universal Reference equimolar pool of 962 synthetic sequences corresponding to human, rat, mouse, and virus miRNA.

Each library preparation was performed by the same two lab scientists using the same equipment. Each protocol was followed exactly as provided by the vendor for each kit. The number of PCR cycles for each sample was determined based on the recommendations of each kit for the various starting input amounts (**Table 2**).

**Table 2.** Number of PCR cycles used for each sample.

| Sample time | Clontech | Illumina | NEB | NEXTflex |
| --- | --- | --- | --- | --- |
| 100ng Brain total RNA | 13 cycles | NA | 15 cycles | 18 cycles |
| 250ng Brain total RNA | 13 cycles | NA | 15 cycles | 18 cycles |
| 500ng Brain total RNA | 11 cycles | NA | 15 cycles | 18 cycles |
| 1000ng Brain total RNA | 10 cycles | 11 cycles | 15 cycles | 18 cycles |
| 1500ng Brain total RNA | 9 cycles | 11 cycles | NA | 18 cycles |
| 2000ng Brain total RNA | 7 cycles | 11 cycles | NA | 18 cycles |
| 300ng of equimolar synthetic pool | 7 cycles | 11 cycles | 15 cycles | 18 cycles |

**Table 2 Legend**. Both batches of 1000ng brain total RNA samples had the same number of cycles. We used the same number of cycles as suggested by each protocol for a range of input amounts. The brain samples were all derived from the same RNA extraction, purchased from Ambion of a 74-year-old Caucasian Female. The cause of death of this individual was respiratory failure. The Miltenyi Biotec miRXplore Universal Reference equimolar pool was used for the accuracy assessments.

Size selection using PAGE gels was recommended by three of the manufacturers (Illumina, NEXTflex, and NEB kits) and was performed for these kits for better comparisons. We used AMPure XP beads for size selection for the Clontech samples, as the vendor does not recommend PAGE gel size selection.

A Qubit Fluorometer was used to determine the concentration of the final libraries. The library preparations were sequenced using single-end sequencing on the Illumina HiSeq 3000 with the Illumina Real Time Analysis (RTA) module and the bcl2fastq2 v2.17 to generate 51 base pair reads.

### Unique Molecular Identifier deduplication

In order to test the use of UMIs to mitigate reverse transcription and amplification bias, reads derived from the NEXTflex kit were collapsed based on random sequences of 4 bases in length contained within both the 3’ and 5’ adapter sequences as a UMI using UMI-tools (49) (**Fig. 1.c**). In this method, the adapters are ligated before reverse transcription and PCR amplification, therefore allowing for the estimation of the abundance of the sequences present in the sample before these steps. In the collapsed NEXTflex data referred to as “Deduped”, only reads that contained the same pair of a unique transcript with a unique UMI were maintained, while duplicate pairs were discarded. Therefore, each unique sequence had the opportunity to bind up to 65,536 different UMIs. As a control, we compared the performance of the Deduped data to a random 5% subset of the reads, referred to as “Fivepercent”. This was necessary as only 5% of the total reads remained following the collapsing method which required a preliminary alignment step. Thus this data was also produced with the preliminary alignment step, all preprocessing was the same except for the use of UMI-tools (49).

We utilized an in-house script to extract the degenerate bases from the adapters to determine the UMI sequence for each read and to add it to the identifier line of the FASTQ files for each sample. In this script we also removed reads which contained any unknown bases within the UMI. We then used *bowtie* (39) (v1.2.2) with a seed length of 15 allowing for 2 mismatches to produce a liberally aligned bam file to be used with UMI-tools (49) for deduping in order to retain miRNA isoforms. With this liberal alignment, various isoforms are aligned to reference miRNAs and thus if any are paired with the same UMI as the reference or other isoforms, they are considered as a duplicate, thus greatly reducing the concern brought up by Sena et al., 2018 (36) about similar inserts being considered different sequences despite pairing with the same UMI instead of being collapsed together. We utilized the directional method in UMI-tools to remove duplicate reads from the bam file, which is particularly stringent for correcting sequencing errors in the UMI sequences themselves (49). We then converted the bam file to a FASTQ file for alignment with miRge (50) with the other method samples. Our script to prepare NEXTflex samples for UMI-tools (49) is available on GitHub at https://github.com/LieberInstitute/miRNA_Kit_Comparison.

### Adapter and degenerate base trimming and alignment

For the NEXTflex (and therefore the Deduped and Fivepercent), NEB, and Illumina FASTQ files the 3’ adapter sequences and all bases 3’ of the adapter were trimmed from the ends of the reads using cutadapt (51) version 1.8.3. For the NEXTflex samples the first and last four bases, which correspond to the random bases included in the adapter sequences, were also trimmed. In the case of the Deduped samples these sequences were added to the identifier line prior to trimming. These bases correspond to the random adapter sequences because sequencing begins at the location of the 4 random bases in the 5’ adapter for this kit.

Unlike the other kits, the Clontech kit is stranded. Read 1 corresponds to the sense strand of the input RNA and the first three bases correspond to the nucleotides added during the SMART template-switching method. Then 10 Adenine bases were removed from the 3’ end, as well as all potential bases 3’ of this stretch of bases.

When trimming the synthetic sample FASTQ files, a lower length limit of 16 bases was used (as this was the shortest synthetic RNA), while a lower length limit of 18 bases was used for the brain samples (as human miRNAs are generally longer than 16 bases), to reduce the inclusion of reads that were too short in the data.

After trimming (and deduping in the case of the Deduped method) samples were aligned to the miRBase human miRNA sequences (52) and the Miltenyi synthetic sequences using miRge (50).

### Similarity Analysis

To perform the hierarchical cluster analysis, we used the miRNA quantifications from all brain libraries with all starting amounts using both batches (total of 99 libraries, 19 for the Clontech, NEXTflex, Deduped and Fivepercent methods and 10 for Illumina and 13 for NEB). We normalized the data using the DESeq2(40) method with the method as the design, using the DESeqDataSetFromMatrix(), estimateSizeFactors(), and counts() functions of the Bioconductor package DESeq2 (40)(v 1.18.1). The DESeq2(40) method was chosen for normalization as we assume little difference between the individual synthetic miRNAs, the replicates across batches, and the triplicates within a given batch given that the samples are biologically the same. Normalization for small RNA sequencing is a debated topic and further studies are needed to confirm the best method for different small RNA sources. We then determined which normalized expression estimates were greater than one for all 99 samples. This resulted in 151 common miRNAs above the threshold. We then log_2_ transformed these estimates. We determined the distance between the samples using the hclust() function of the stats package (v 3.4.0). We also used these quantification estimates in a sum of squares analysis to determine the percent of variance explained by method, starting amount, batch, and the number of reads that mapped to miRNA. To do this we used the Anova() type II function of the CRAN car package(v 3.0-0). To create the MA plots we used only the 1000ng brain samples from both batches (a total of 24 samples, 4 for each method). We normalized this subset of samples using again using DESeq2(40) and method as the design. We again restricted our analysis to miRNAs with greater than one normalized count in all 24 samples. This resulted in 174 common miRNAs above the threshold. We then manually created the MA plots. We then ranked the log2 normalized quantifications and determined the overlap of the most abundant miRNAs.

### Accuracy Assessment

To perform the accuracy analyses, we used equimolar pools of 962 synthetic miRNAs purchased from Miltenyi Biotec. The Gibbs minimum free energy of the secondary structures for each synthetic miRNA was determined using RNAfold as part of the ViennaRNA package 2 (version 2.3.5) (53, 54) GC content was determined using the letterFrequency() function of the Bioconductor package Biostrings (55) (v 2.46.0). Alignments were performed using the miRge program. The miRge raw count estimates were normalized using DESeq2 (40) (v 1.18.1) with method as the design. The difference of each miRNA count estimate was then calculated from the mean of all synthetic sequences. The absolute of this difference was then log_2_ transformed for statistical comparisons and is referred to as “accuracy error”.

A linear model was used to evaluate the influence of method on accuracy error, using the lm() function of the stats package, and paired t-tests using the t.test() function of the stats package (v 3.4.0) were used for pairwise comparisons of each method. The Bonferroni method was used to for multiple testing correction. The omega squared value was calculated using the anova_stats() function of the CRAN package sjstats (56) (v 0.16.0). Hedge’s g was calculated using the tes() function of the CRAN package compute.es (v 0.2-4). Catplots to evaluate concordance of error rank were created using the CATplot() function of the Bioconductor package ffpe (v 1.22.0).

### Detection Diversity Assessment

To assess mapping rates to various classes of RNAs, we collected FASTA files for a variety of RNAs: miRNA, piRNA, rRNA, scaRNA, snoRNA, snRNA, and transfer RNA (tRNA) and then created a merged FASTA file from each of the smaller FASTA files. We used the miRNA FASTA file included in miRge. The piRNA data was acquired from piRNAQuest (57) (http://bicresources.jcbose.ac.in/zhumur/pirnaquest/). The rRNA, tRNA, and snRNA data came from the hg19 assembly from the UCSC genome table browser (58) (http://genome.ucsc.edu). The snoRNA and scaRNA data came from snoRNABase (59) (https://www-snorna.biotoul.fr/browse.php). Only the C/D box snoRNAs were included as all the H/ACA box snoRNAs overlapped with the snRNA data from UCSC. Six of the C/D snoRNA sequences and the snRNA overlapped in our merged FASTA file. Additionally, all of the scaRNAs overlapped the snoRNA C/D box sequences, but we maintained them in order to analyze scaRNA. Exact matches of miRNA sequences and piRNA were removed from the piRNA portion of the FASTA file. *bowtie* (39) was used for alignment to all the sequences simultaneously allowing for zero mismatches within the default seed length of 28 bases to better distinguish similar sequences of different RNA classes. We then determined the count of reads that mapped to each RNA class.

miRNAs were considered detected if they were observed with > 10 normalized reads in all triplicates of a given starting amount. Raw counts from miRge for the brain batch 1 samples (93 in total) were normalized with DESeq2(40) but were not log transformed. Another analysis was performed using both batches and normalizing with DESeq2(40) using all brain samples and a threshold of >10 normalized reads for all samples of a given starting amount. The percent of detected miRNAs that were inconsistently detected was calculated as follows:

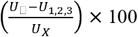

Where *U_x_* = number of unique miRNAs detected with >10 normalized reads in a given triplicate and where *U*_1,2,3_ = number of unique miRNAs detected with >10 normalized reads in all triplicates.

The same methods were used for the isomiR analysis, however a threshold of 100 normalized reads was used instead of 10.

Statistical analyses of the detection and detection inconsistency were performed as in the Accuracy assessment with lm() and t.test() of stats package (v 3.4.0) and the tes() function of the compute.es package (v 0.2-4) and the anova_stats() function of the sjstats package (v 0.16.0) to calculate effect sizes. Percent variance explained analyses of sequence characteristics were performed using the Anova() function of the CRAN package car (v 3.0-0). Concordance was evaluated using the CATplot() function of the Bioconductor package ffpe (v 1.22.0). The evaluation of false isomiRs used the synthetic miRNA data. An isomiR was considered as detected if over 100 normalized reads were observed.

### Consistency Assessment

Consistency across batch was determined using all 1000 ng brain samples (24 samples). DESeq2(40) (v 1.18.1) was used to normalize these samples with method as the design. Quantifications were filtered for those with >10 normalized reads in all samples of a given method. The mean of the quantifications from the first batch triplicates was compared with that of the second batch quantifications. The log_2_ transformed value of the absolute difference between these two quantifications was used to compare the batch consistency of the methods. Again the lm() of the stats package (v 3.4.0) was used for global analyses, while the t.test() function with Bonferroni correction was used to compare pairs of methods. Evaluating the intersection of all miRNAs detected across both batches for each method (total of 162 miRNAs), we determined the percent of variance in triplicate error for sequence characteristics.

To evaluate the consistency of triplicates, we used all 93 brain samples of the first batch. This data was normalized using DESeq2(40) using method as the design. The quantification estimates were filtered for those with >10 normalized counts in all samples for a given starting amount. “Triplicate error” was determined as the difference of the value of each triplicate relative to the mean of all triplicates. The absolute value of these differences was then log_2_ transformed and the mean error value of triplicates was determined for each miRNA detected by each method for statistical comparisons. Evaluating just the intersection of all miRNAs detected for each starting amount and method (total of 228 miRNAs), we determined the percent of variance in triplicate error for sequence characteristics. The consistency of triplicates was then used to compare the various starting amounts. The Bonferroni method was used for multiple testing correction.

## Computational Resources

The code for all of the analyses performed in this manuscript is will be publically available at https://github.com/LieberInstitute/miRNA_Kit_Comparison. The data will be made available at the National Institutes of Health (NIH) Sequence Read Archive (SRA), and the accession number will be listed on the GitHub readme for the repository.

## Author’s contributions

C.W. (Carrie Wright) designed the study, performed the data analysis, and wrote the manuscript. C. W. (Carrie Wright) and A.R. performed the library preparations and data preprocessing. A.R. also edited the manuscript and assisted with the study design and the design for **Fig. 1**. A.R. wrote the script to prepare the FASTQ files for UMI-tools with the assistance of C.W. (Carrie Wright). E.E.B. assisted with creating the figures and edited the manuscript. C.W. (Courtney Williams) and M.K. performed the sequencing. L.C.-T. and A.E.J assisted with the statistical analysis design. L.C.-T, A.E.J, D.R.W., and J.H.S. also edited the manuscript. J.H.S. supervised this project and assisted with the study design and the design of **Fig. 1**. A.J.C., N.J.B., and D. R.W. secured funding for this project and contributed to the overall direction.

## Acknowledgements

We are grateful for the generosity of the Lieber and Maltz families in establishing an institute dedicated to understanding the basis of developmental brain disorders. We are grateful to the Johns Hopkins Program for microRNA Biology Journal Club for feedback about our study, particularly from Marc Halushka.

## Supplementary FIGURE LEGENDS

**Supplementary Fig. 1**. Boxplots and heatmaps of the influence of various synthetic sequence aspects on accuracy error.

Boxplots show sequences grouped by various sequence aspects, including: Gibb’s free energy secondary structure estimates, the identity of the last 2 bases, the identity of the first 2 bases, and the number of Cs in the sequence. The percent expression relative to the mean of all synthetic sequences is plotted for each of the 962 synthetic sequences. Heatmaps also depict the percent expression of synthetic sequences grouped by the sequence aspect of interest relative to the mean expression of all the synthetic sequences. Overall, most methods showed a positive quantification relationship with higher or less negative Gibb’s free energy estimates. Most methods showed consistent quantification despite the identity of the last two bases, however NEB and Illumina showed more inconsistent quantification. All of the methods showed fairly consistent quantification despite the identity of the first two bases, however the Deduped method showed more consistent quantification. Most of the methods showed decreased quantification with increasing numbers of Cs, however the Clontech method showed the opposite relationship, and the Deduped method was quite consistent.

**Supplementary Fig. 2**. Detection of miRNAs and isomiRs and detection consistency of miRNA and isomiRs across various starting amounts.

For the detection plots, the number of miRNAs detected above 10 normalized counts and the number of isomiRs detected above 100 normalized counts in all triplicates of batch 1 for each method is plotted on the y-axis. The starting total RNA amount is indicated on the x-axis in nanograms. Only starting amounts in the range of suggested inputs were tested for each method. For the consistency of detection plots, the percentage of miRNAs or isomiRs detected above the threshold by a single triplicate that were not detected above the threshold by the other two triplicates is plotted on the y-axis. The starting total RNA amount is indicated on the x-axis in nanograms. The relationship between the percentage of inconsistently detected miRNAs or isomiRs and starting amount is plotted as a line using a locally estimated scatterplot smoothing regression (LOESS).

