## Supplementary Figure 1 for "Comprehensive assessment of multiple biases in small RNA sequencing reveals significant differences in the performance of widely used methods"

### Influence of estimated secondary structure on accuracy error

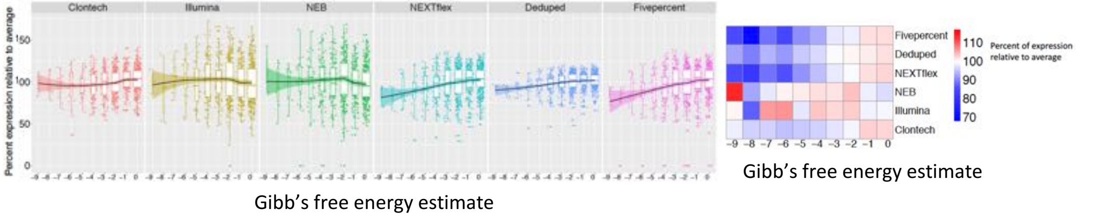


### Influence of the identity of the last 2 bases in synthetic sequences on accuracy error

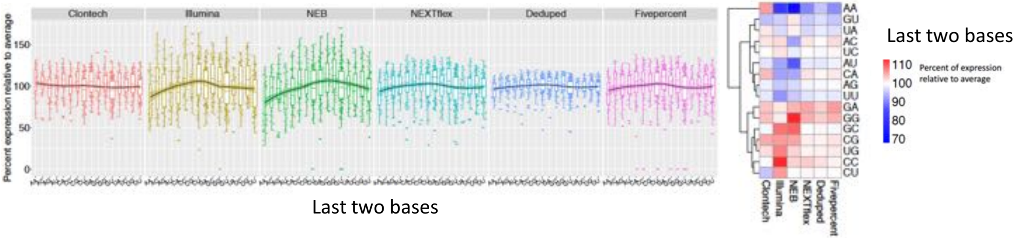


### Influence of the identity of the first 2 bases in synthetic sequences on accuracy error

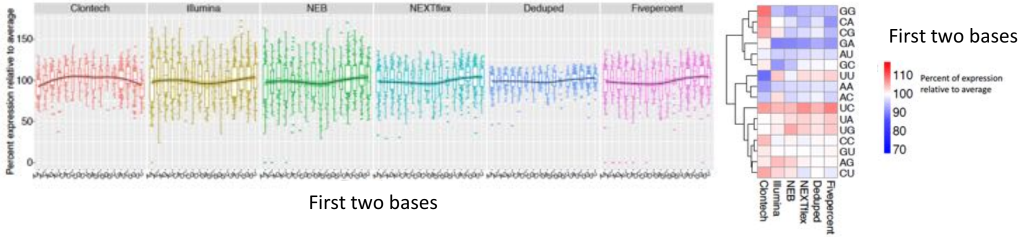


### Influence of the number of Cs in synthetic sequences on accuracy error

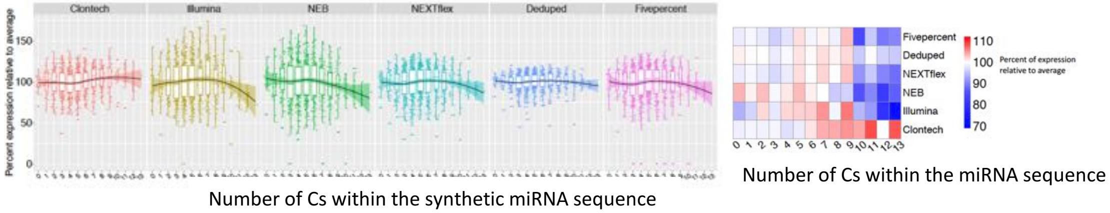
