## Supplementary Figure 2 for "Comprehensive assessment of multiple biases in small RNA sequencing reveals significant differences in the performance of widely used methods"

**miRNA Detection across kits > 10 reads in all triplicates**

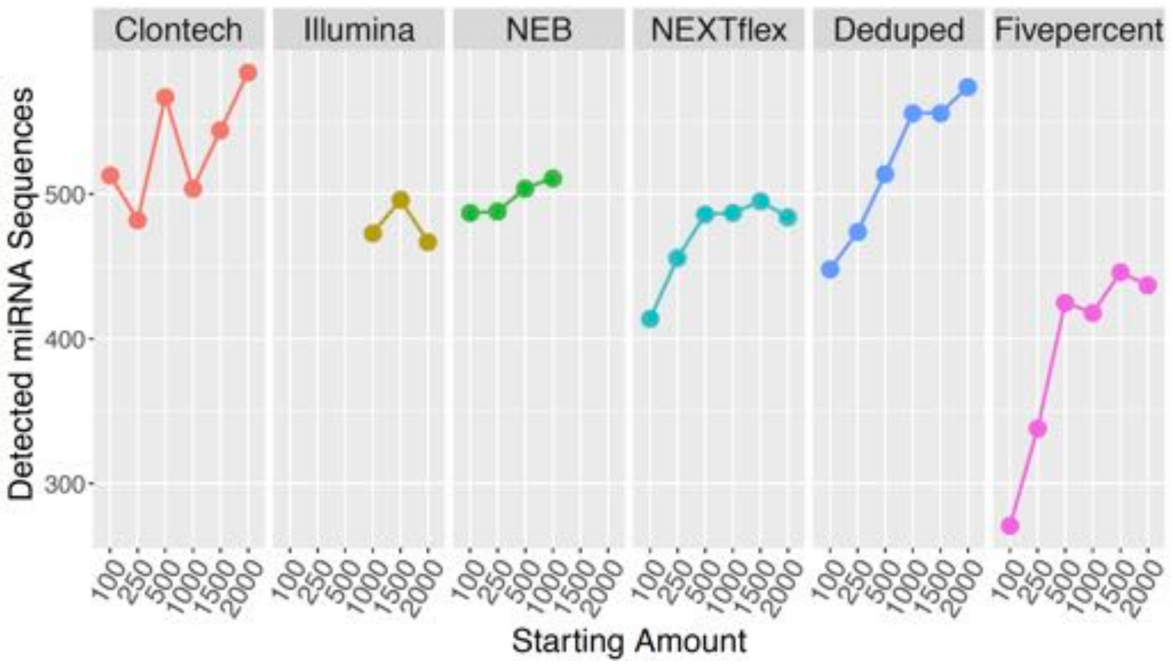


**isomiR Detection across kits > 100 reads in all triplicates**

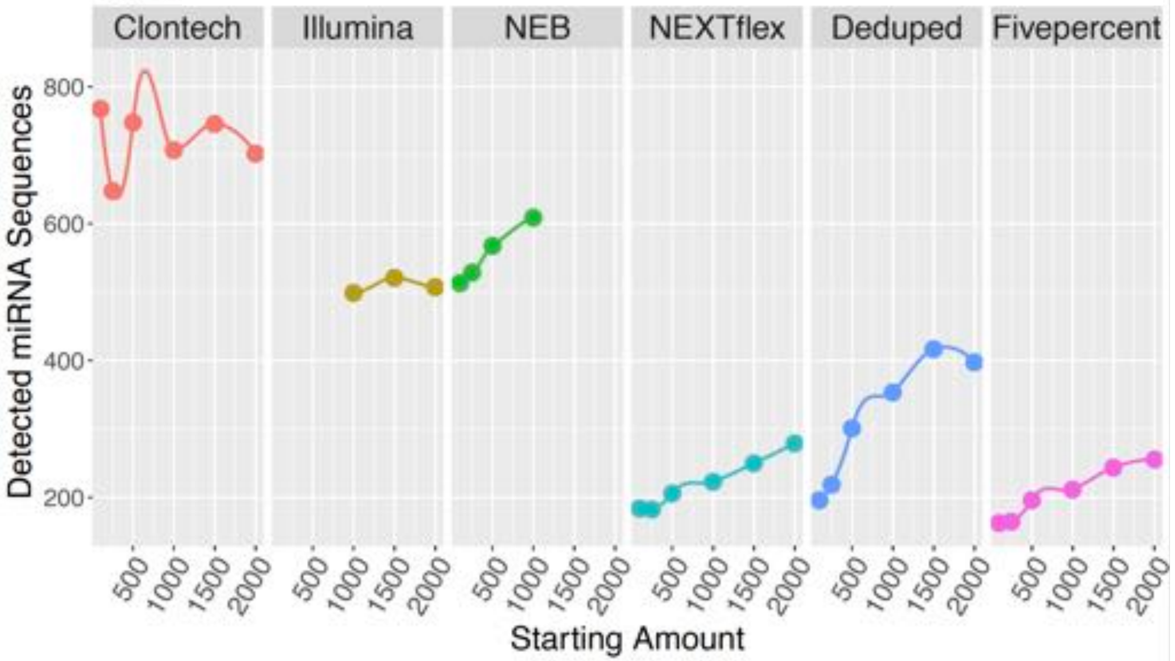


**Inconsistency of miRNA detection among triplicates**

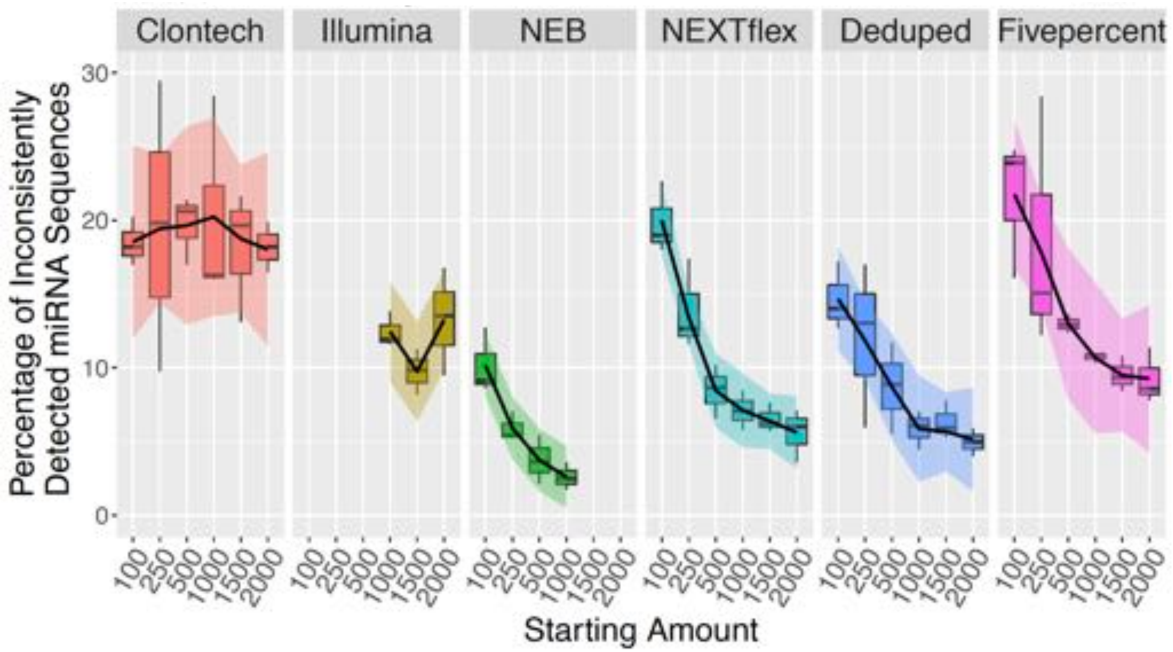


**Inconsistency of isomiR detection among triplicates**

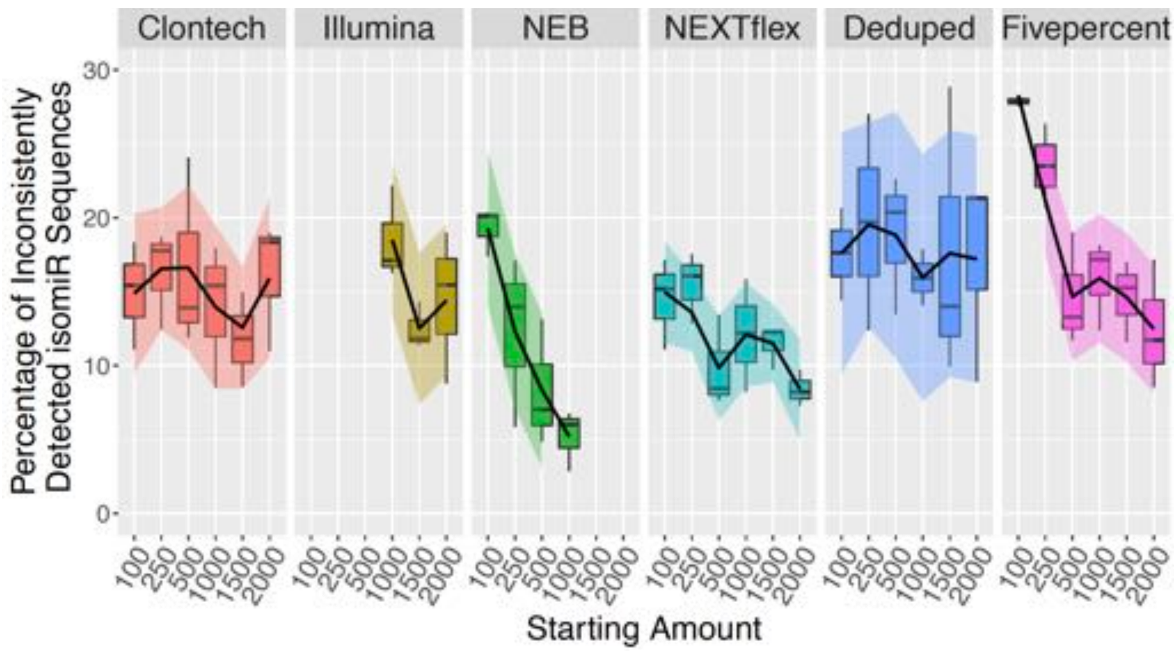
