## Supplementary Note 1 for "Comprehensive assessment of multiple biases in small RNA sequencing reveals significant differences in the performance of widely used methods"

With the 4 random bases in each adapter sequence, our UMIs were 8 bases long. With four different possible bases this results in $4^8$ or 65,536 possible unique UMI sequences. Therefore, each unique miRNA sequence present before reverse transcription and amplification had ideally 65,536 possible UMI sequences to bind.

The question is then, how often a unique miRNA sequence is present with more than 65,536 copies in a given sample before amplification. With longer UMI sequences a larger number of copies could be accommodated. It is important to note that often miRNA counts include reads that map to a variety of isomiR sequences. In this case, multiple sequences would each have 65,536 different UMI sequences to bind.

To determine if our UMI length was adequate, we evaluated the number of reads mapping to the most abundant miRNA among the NEXTflex samples, miR-9-5p. This miRNA accounted for at most 13% of all miRNA mapping reads, when including all isoforms, and at most 11.6% when including only the canonical isoform. Back-calculating an estimate of the number of canonical miR-9-5p sequences present in the samples before amplification, we determined that all of the samples regardless of starting input had fewer than the number of unique UMI sequences (65,536). For 18 PCR cycles, we calculated this estimate as:

$$\frac{A}{E^{18}}$$

Where $A$ = Number of reads mapping the most abundant isoform which was the canonical in this case, $E^N$ = the PCR efficiency rate to the power of the $N$ number of PCR cycles, in this case 18.

This remained true even when assuming a PCR efficiency as low as 62.5% (where $E$ = 1.25, as 1.25/2 = 62.5% and perfect efficiency would double all molecules at each cycle). We calculated this estimate as:

$$\frac{A}{1.25^{18}}$$

Assuming this low level of efficiency, an average of 21,060 miR-9-5p sequences were estimated as initially present, with only one sample approaching the upper limit of the unique UMI sequences, with an estimate of 63,672 initial sequences. Previous studies suggest that miRNA PCR efficiency is generally in the range of 75 to 95% [1], and a previous miR-9-5p PCR study showed efficiencies of roughly 98% [2]. Therefore, this suggests that our UMI was likely long enough to accommodate all miRNAs assessed in this study. However, in tissues that may have more predominant small RNAs or in studies with greater sequencing depth, a longer UMI may be required. See **Supplementary Table 24** for the estimates of starting molecules for each NEXTflex samples for miR-9-5p and a less abundant miRNA, miR-137.

1. Brunet-Vega, A. et al. Data on individual PCR efficiency values as quality control for circulating miRNAs. Data Brief 5, 321–326 (2015).

2. Barbano, R. et al. Stepwise analysis of MIR9 loci identifies miR-9-5p to be involved in Oestrogen regulated pathways in breast cancer patients. Sci. Rep. 7, (2017).
